# The biomechanical role of extra-axonemal structures in shaping the flagellar beat of Euglena

**DOI:** 10.1101/2020.03.15.991331

**Authors:** Giancarlo Cicconofri, Giovanni Noselli, Antonio DeSimone

**Affiliations:** SISSA - International School for Advanced Studies - Trieste (Italy); The BioRobotics Institute, Scuola Superiore Sant’Anna - Pisa (Italy)

## Abstract

We propose and discuss a model for flagellar mechanics in *Euglena gracilis*. We show that the peculiar non-planar shapes of its beating flagellum, dubbed “spinning lasso”, arise from the mechanical interactions between two of its inner components, namely, the axoneme and the paraflagellar rod. The spontaneous shape of the axoneme and the resting shape of the paraflagellar rod are incompatible. The complex non-planar configurations of the coupled system emerge as the energetically optimal compromise between the two antagonistic components. The model is able to reproduce the experimentally observed flagellar beats and their characteristic spinning lasso geometric signature, namely, travelling waves of torsion with alternating sing along the length of the flagellum.

## Introduction

Flagella and cilia propel swimming eukaryotic cells and drive fluids on epithelial tissues of higher organisms [1]. The inner structure of the eukaryotic flagellum is an arrangement of microtubules (MTs) and accessory proteins called the axoneme (Ax). A highly conserved structure in evolution, the Ax typically consists of nine cylindrically arranged MT doublets cross-bridged by motor proteins of the dynein family. An internal central pair of MTs is connected by radial spokes to the nine peripheral doublets, determining the typical “9+2” axonemal structure. Motor proteins hydrolyze ATP to generate forces that induce doublet sliding. Due to mechanical constraints exerted by linking proteins (nexins) and the basal body, dynein-induced sliding of MTs translates into bending movements of the whole structure. Motor proteins are thought to self regulate their activity via mechanical feedback, generating the periodic beats of flagella, see e.g. [6] and [19].

Despite a general consensus on the existence of a self-regulatory mechanism, the inner working of the Ax is still not fully understood and it is still a subject of active research [33]. While bending-through-sliding is the accepted fundamental mechanism of flagellar motility, how specific flagellar shapes are determined is not yet clear. For the most studied swimming microorganisms, such as animal sperm cells and the biflagellate alga *Chlamydomonas reinhardtii*, the flagellar beat is, to a good approximation, planar. For these organisms, beat planarity is thought to be induced by the inter-doublet links between one pair of MTs, typically those numbered 5 and 6 [18]. These links inhibit the relative sliding of the 5-6 MTs pair, thus selecting a beating plane that passes through the center of the Ax and the midpoint between the inhibited MTs.

A remarkable deviation from the flagellar structure of the aforementioned organisms is found in euglenids and kinetoplastids. These flagellated protists have a whole extra element attached alongside the Ax [7], a slender structure made of a lattice-like arrangement of proteins called “paraxial” or “paraflagellar” rod (PFR), see Figure 1. The latter name is more common, but the former is possibly more accurate [24]. PFRs are attached via bonding links to up to four axonemal MTs, depending on the species [32]. PFRs are thought to be passive but, at least in the case of Euglena, some degree of activity is not completely ruled out [21].

**Figure 1.**
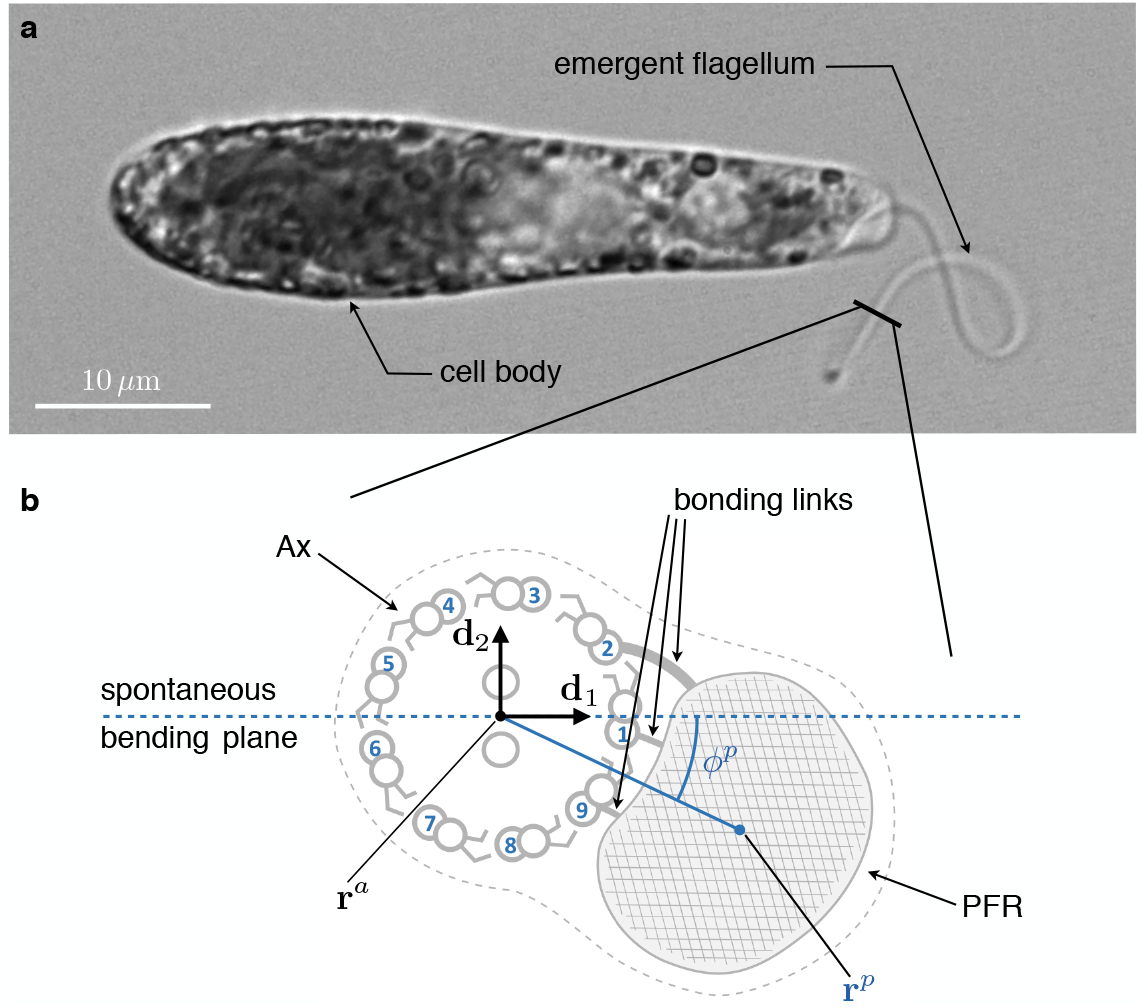
a) A specimen of freely swimming *Euglena gracilis*, and b) a sketch of the cross section of its flagellum seen from the distal end. The flagellar inner structure is composed by the paraflagellar rod (PFR, textured), and the axoneme (Ax). The PFR is connected via bonding links to the axonemal doublets 1, 2, and 9 (our numbering convention). The inner structure of the flagellum is enclosed by the flagellar membrane (dotted line). By inhibiting MTs’ sliding the PFR selects the spontaneous bending plane of the Ax (dashed line). The solid line that joins the Ax center **r**^*a*^ and the PFR center **r**^*p*^ cross at an angle *ϕ*^*p*^ the spontaneous bending plane.

In this paper, we put forward and test the hypothesis that the distinctive beating style of *Euglena Gracilis*, sometimes dubbed “spinning lasso” [5], arises due to PFR-Ax mechanical interaction.

In order to put our hypothesis into context, we observe that the flagellar beat of PFR-bearing kinetoplastid organisms, such as Leishmania and Crithidia, is planar [9]. An apparent exception to beat planarity in kinetoplastids is found in the pathogenic parasite *Trypanosoma brucei*, which shows a characteristic non-planar “drill-like” motion [17]. However, Trypanosoma’s flagellum is not free, like that of Leishmania and Crithidia, but it is attached to the organism for most of its length, wrapped helically around the cell body. According to [2] the flagellum-body mechanical interaction could alone explain Trypanosomes’ distinctive motion. Confirming this conclusion, [34] showed that Trypanosoma mutants with body-detached flagellum generate fairly planar beating. It is conjectured that the PFR-Ax bonds operate as the 5-6 interdoublet links in Chlamydomonas and sperm cells, inhibiting MTs sliding and selecting a plane of beat [36].

Euglena’s spinning lasso beat does not conform to this scenario. Indeed, the beating style of Euglena is characterized by high asymmetry and non-planarity. The full 3d flagellar kinematics of freely swimming cells has recently been revealed [25] thanks to a mixed approach based on hydrodynamic theory and image analysis. As we report in the first part of this paper, the geometry of the spinning lasso is characterized by travelling waves of torsion with alternating sign along the flagellum length.

We argue that the key to the emergence of non-planarity lies in a prominent asymmetry in the structure of PFR-Ax attachment in euglenid flagella. Figure 1 shows a sketch of the cross section of the euglenid flagellum redrawn from [20], see also [4]. The PFR is attached to three MTs, which we number 1, 2, and 9. We consider two lines. One line (dashed) passes through the center of the Ax and MT 1, in the middle of the bonding complex. The other line (solid) connects the center of the Ax and the center of the PFR. The two lines cross each other. This is the structural feature on which we build our model.

In the model we assume that the bonding links to the PFR select the local spontaneous beating plane of the Ax, from the same principle of MTs’ sliding inhibition discussed above. The local spontaneous beating plane so generated passes through the dashed line in Figure 1. We follow closely [14] and [27] in our modeling of the Ax, while we use a simple elastic spring model for the PFR. We show that, under generic actuation, the two flagellar components cannot be simultaneously in their respective states of minimal energy, and this crucially depends on the offset between the spontaneous beating plane of the Ax (dashed line in Figure 1) and the line joining the PFR-Ax centers (solid line in Figure 1). Instead, the typical outcome is an elastically frustrated configuration of the system, in which the two competing components drive each other out of plane. Under dyneins activation patterns that, in absence of extra-axonemal structures, would produce an asymmetric beat similar to those of Chlamydomonas [23], or Volvox [26], the model specifically predicts the torsional signature of the spinning lasso, which we discuss in the following Section.

Interestingly, the lack of symmetry of the spinning lasso beat produces swimming trajectories with rotations coupled with translations [25]. In turn, cell body rotations have a role in phototaxis (see Discussion and Outlook Section and [12]). In the light of these observations, our analysis shows that the beat of the euglenid flagellum can be seen as an example of a biological function arising from the competition between antagonistic structural components. It is not dissimilar from the body-flagellum interaction in Trypanosoma, which generates 3d motility. But the principle is much more general in biology and many other examples can be found across kingdoms and species and at widely different scales. For instance in plants, a mechanism of seed dispersal arises from the mechanical competition between the two valves of the seed pods, see, e.g., [3] for *Bauhinia variegata* and [15] for *Cardamine hirsuta*. Contraction by antagonistic muscles is key for animal movement and, in particular, for the functioning of hydrostatic skeletons (used from wormlike invertebrates to arms and tentacles of cephalopods, to the trunk of elephants, see [16]). A similar principle of antagonistic contraction along perpendicularly oriented families of fibers is at work at the sub-cellualr level, for example in the antagonistic motor protein dynamics in contractile ring structures important in cell division and development (see, e.g., [8]). At the same sub-cellular scale, competing elastic forces arising from lipid-protein interactions are often crucial in determining the stability of complex shapes of the cellular membrane [31], and in the case of the overall structure of the coronavirus envelope [29].

## Observations

We first analyze the experimental data from the 3d reconstruction of the beating euglenid flagellum obtained in [25] for freely swimming organisms. Swimming Euglenas follow generalized helical trajectories coupled with rotation around the major axis of the cell body. It is precisely this rotation that allows for a 3d reconstruction of flagellar shapes from 2d videomicroscopy images. Euglenas take many beats to close one complete turn around their major body axis. So, while rotating, Euglenas show their flagellar beat to the observer from many different sides. Stereomatching techniques can then be employed to reconstruct the flagellar beat in full (assuming periodicity and regularity of the beat).

Figure 2 shows *N* = 10 different curves in space describing the euglenid flagellum in different instants within a beat taken from [25]. The reconstruction fits well experimental data from multiple specimens. The figure also illustrates the calculated torsion of the flagellar curve at each instant (not previously published). Torsion, the rate of change of the binormal vector, is the geometric quantity that measures the deviation of a curve from a planar path (see the Results Section below for the formal definition). The spinning lasso shows here a distinct torsional signature characterized by torsion peaks of alternate sign, traveling from the proximal to the distal end of the flagellum.

**Figure 2.**
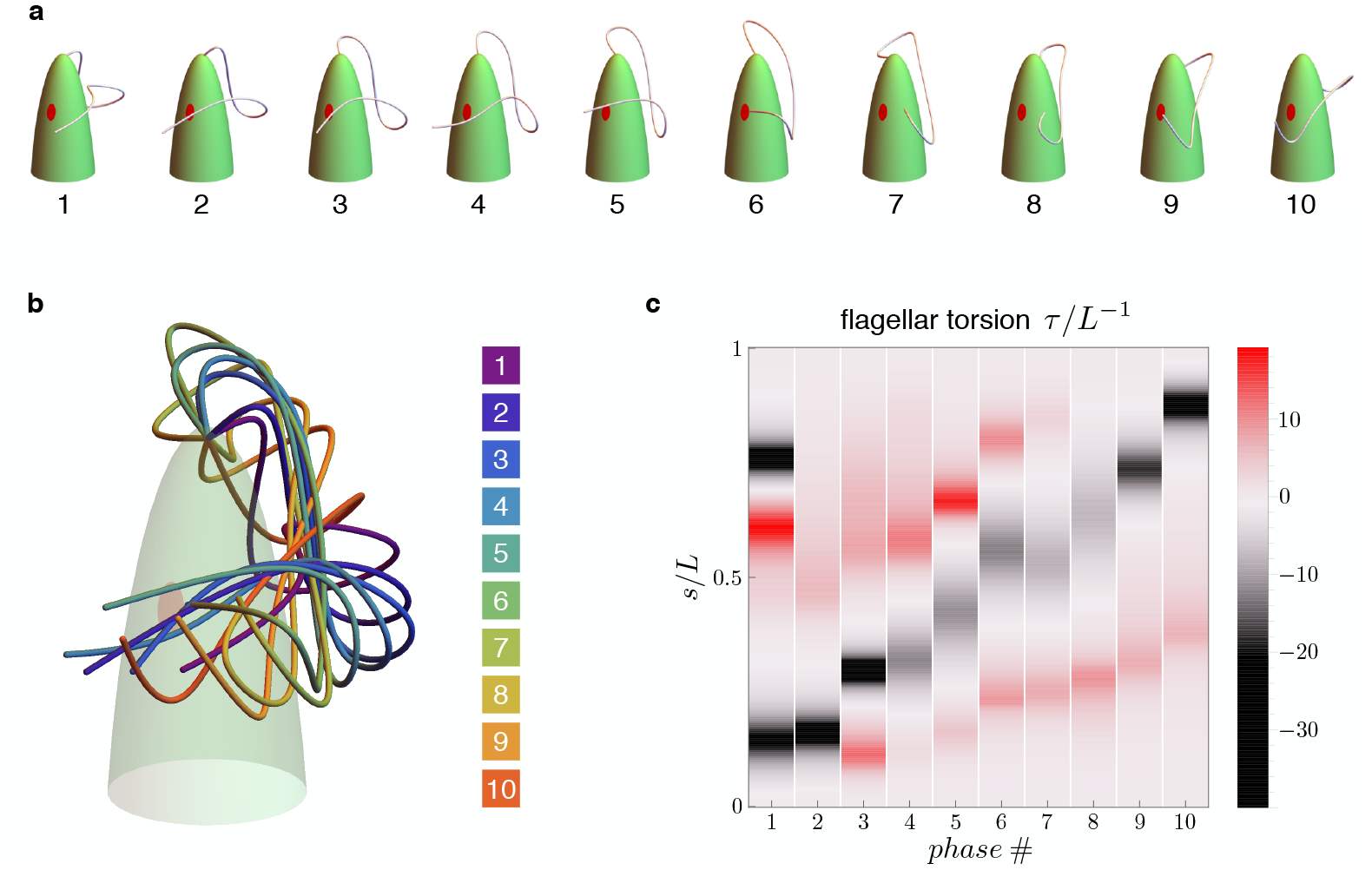
Flagellar beat kinematics of freely swimming *Euglena gracilis*. a) *N* = 10 flagellar configurations in evenly spaced instants (phases) within a periodic beat. b) The same configurations overlapped and color coded according to their phases. c) Calculated torsion *τ* = *τ* (*s*) as a function of the flagellar arc length *s*. The plot is presented in terms of the normalized quantities *τ/L*^−1^ and *s/L*, where *L* is the total length of the flagellum.

To further investigate Euglena’s flagellar beat we observed stationary cells trapped at the tip of a capillary. In this setting the flagellum is not perturbed by the hydrodynamic forces associated with Euglena’s rototranslating swimming motion. The beat can then manifest itself in its most “pristine” form. We recorded trapped Euglenas during periodic beating. Then, we rotated the capillary and recorded the same beating cell from different viewpoints. Videomicroscopy images from one specimen are shown in Figure 3. While with fixed specimens we cannot reconstruct reliably the 3d flagellar shapes, Figure 3 shows that there is a high stereographical consistency with the flagellar shapes obtained from swimming organisms. Flagellar non-planarity is thus not intrinsically associated with swimming, which reinforce the idea that the mechanism that generates non-planar flagellar shapes might be structural in origin. Moreover, these observations justify the choice we made in our study to focus on a model of flagellar mechanics for stationary organisms, allowing for substantial simplifications.

**Figure 3.**
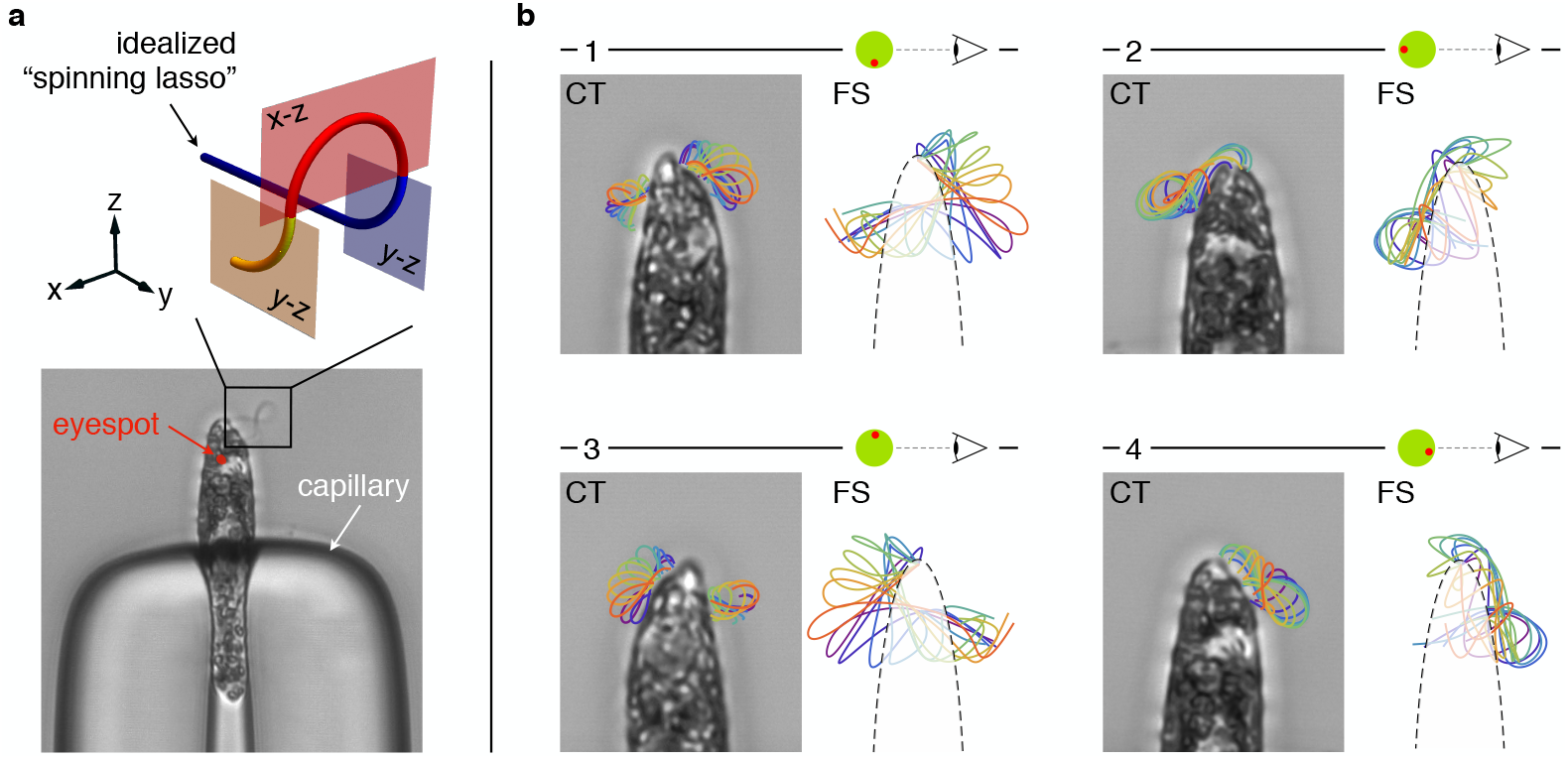
a) A specimen of *Euglena gracilis* trapped at the tip of a capillary (bottom). The typical outline of its beating flagellum is that of a looping curve, which is consistent with the outline of a simple curve with two concentrated torsional peaks of alternate sign along the length of the curve, i.e. a torsional dipole (top). b) Close-up images of the same specimen of capillary-trapped (CT) Euglena seen from different viewpoints, upon successive turns of the capillary tube. The body orientation with respect to the objective is estimated from the anatomy of the cell, and in particular from the position of the eyespot (a visible light-sensing organelle present on the cell surface). Microscopy images are decorated with the tracked outlines of the flagellum in different phases (same color coding as in Figure 2). The outlines (2d projections) of the 3d reconstructed flagellar beat of freely swimming (FS) specimens are shown for comparison.

As a final remark we observe how, from a simple geometric construction, we can show that the torsional pattern in Figure 2 is consistent with Euglena’s flagellar shapes as seen from common 2d microscopy, for either swimming or trapped organisms. Typically, the 2d outline (i.e. the projection on the optic plane) of a beating euglenid flagellum is that of a looping curve, see e.g. [30] for independent observations. Consider now an idealized 3d model of the spinning lasso geometry: a curve with two singular points of concentrated torsion with opposite sign, such as the one shown in Figure 3. If we move along the curve, from proximal end to distal end, we first remain on a fixed plane (blue). Then the plane of the curve abruptly rotates by 90 degrees (red plane) first, and then back by 90 degrees in the opposite direction (yellow plane). These abrupt changes correspond to concentrated torsional peaks of opposite sign. When seen in a two dimensional projection, this torsion dipole generates a looping curve that closely matches euglenid flagella’s outlines during a spinning lasso beat.

### Mechanical model

We model Ax and PFR as cylindrical structures with deformable centerlines, see Figure 4. The euglenid flagellum is the composite structure consisting of Ax and PFR attached together. We suppose that the Ax is the only active component of the flagellum, whereas the PFR is purely passive. Our mechanical model builds on the definition of the total internal energy of the flagellum

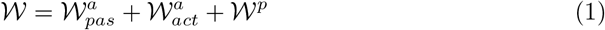

which is given by the sum of three terms: the passive (elastic) internal energy 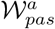 of the Ax, the active internal energy 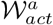 of the Ax (generated by dynein action), and the (passive, elastic) internal energy 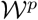 of the PFR. The passive internal energy of the Ax is given by

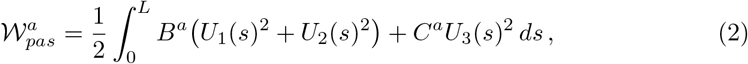

which is formally identical to the classical expression for the energy of elastic (inextensible) rods. We denoted with *U*_1_ and *U*_2_ the bending strains, *U*_3_ is the twist, while *B*^*a*^ and *C*^*a*^ are the bending and twist moduli, respectively. *L* is the total length of the Ax centerline **r**^*a*^. Bending strains and twist depend on the arc length *s* of the centerline, and they are defined as follows. We associate to the curve **r**^*a*^ an orthonormal frame **d**_*i*_(*s*), with *i* = 1, 2, 3, which determines the orientation of the orthogonal sections of the Ax (enclosed by light blue circles in Figure 4). The unit vectors **d**_1_(*s*) and **d**_2_(*s*) define the plane of the orthogonal section at *s*. The unit vector **d**_3_(*s*) = *∂*_*s*_**r**^*a*^(*s*) lies perpendicular to the section. Bending strains and twist are then given by

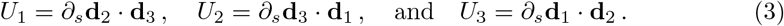

**Figure 4.**
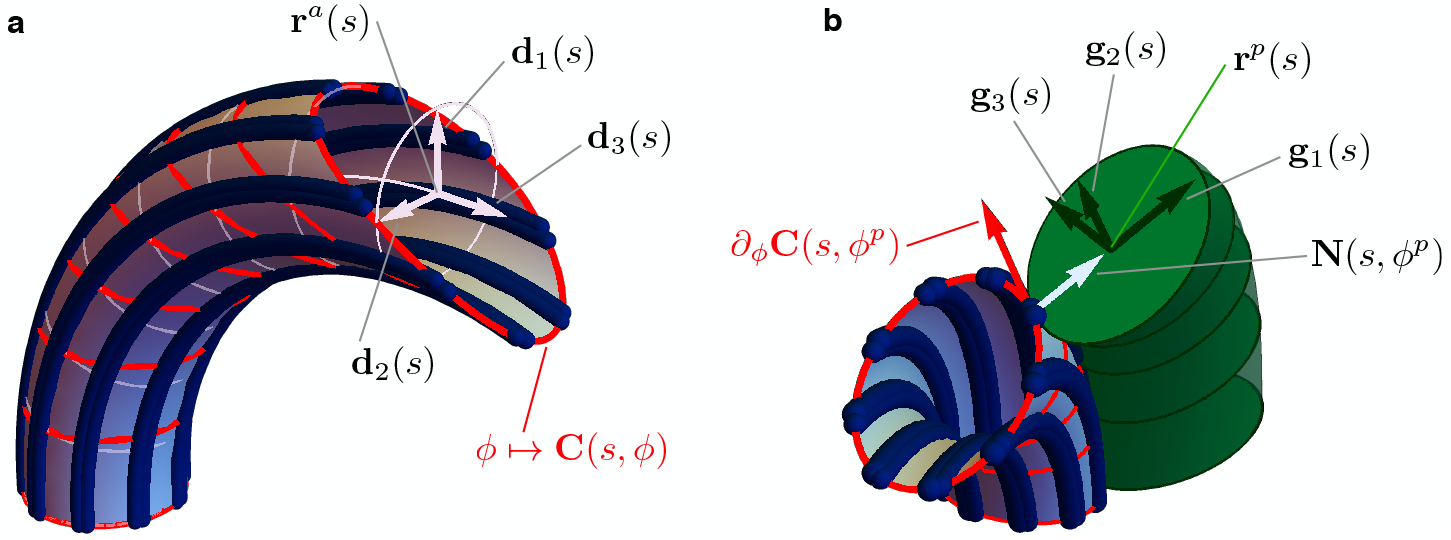
a) Geometry of the Ax. MTs lie on a tubular surface **C**(*s, ϕ*) parametrized by generalized polar coordinates *s* and *ϕ*, where *s* is the arc length of the axonemal centerline **r**^*a*^. The unit vectors **d**_1_(*s*) and **d**_2_(*s*) lie on the orthogonal cross sections of the Ax (light blue circles). The material sections of the Ax are given by the curves *ϕ* ↦ **C**(*s, ϕ*) (red), which connect points of neighbouring axonemal MTs corresponding to the same arc length *s*. Bend deformations of the axoneme are generated by the shear (collective sliding) of MTs. The shear is quantified by the angle between the orthogonal sections and the material sections of the Ax. b) Geometry of the euglenid flagellum, detail of the Ax-PFR attachment. The unit vectors **g**_1_(*s*) and **g**_2_(*s*) generate the plane of the PFR’s cross sections. The vector **g**_1_(*s*) is parallel to the outer unit normal to the axonemal surface **N**(*s, ϕ*^*p*^), while **g**_2_(*s*) is parallel to the tangent vector to the material section *∂*_*ϕ*_**C**(*s, ϕ*^*p*^).

Thus, *U*_1_ and *U*_2_ measure the bending of the Ax on the local planes **d**_1_-**d**_3_ and **d**_2_-**d**_3_, respectively, while the twist *U*_3_ is given by the rotation rate of the orthonormal frame around the centerline’s tangent **d**_3_.

We remark here that we do not consider the Ax as a “filled” beam but as a hollow tubular structure; its elasticity comes from the individual MTs lying on its outer surface. The derivation of (2) from a detailed model of the Ax is given in Appendix A.

The active internal energy of the Ax is defined as minus the total mechanical work of the dyneins

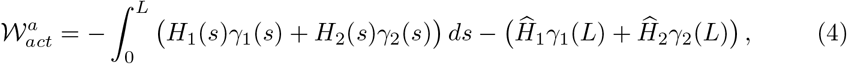

where *γ*_1_ and *γ*_2_ are the two variables that quantifies the shear (i.e. collective sliding) of MTs, while *H*_1_ and *H*_2_ are the corresponding shear forces exerted by molecular motors. Following [28] we also allow for singular shear forces, 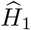 and 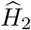, concentrated at the distal end of the Ax. This forces arise naturally, as we remark after (22) in the Results Section. The shear variables are related to the MTs’ kinematics by the following formulas. The MTs’ centerlines **r**^*j*^, for *j* = 1, 2 . . . 9, are given by **r**^*j*^(*s*) = **C**(*s, ϕ*_*j*_), where *ϕ*_*j*_ = 2*π*(*j* − 1)/9, and

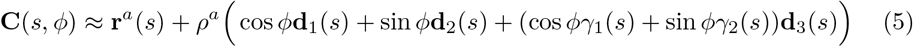

is the parametrization of the cylindrical surface of the Ax (*ρ*^*a*^ is the Ax radius) in terms of the centerline arc length *s* and the angle *ϕ*. In Appendix A we show how the shear variables are related to the individual sliding of MTs with respect to their neighbours (35), and we compute the relations between the dynein forces acting on each pair of adjacent MTs and the shear forces *H*_1_ and *H*_2_ (36). A special axonemal deformation with *γ*_2_ = 0 is shown in Figure 4. The Ax is in this case bent into a circular arc, and the centerline **r**^*a*^ lies on the plane generated by the unit vectors **d**_1_ and **d**_3_. The shear variable *γ*_1_(*s*) ≠ 0 has here a simple geometrical interpretation. For each fixed *s* the curve *ϕ* ↦ **C**(*s, ϕ*) describes what we call the “material” section of the Ax at *s* (red curves in Figure 4). The material section is a planar ellipse centered in **r**^*a*^(*s*) which connects points of neighbouring MTs’ corresponding to the same arc length. Formula (5) says that *γ*_1_(*s*) is the tangent of the angle at which the material sections at *s* intersect the orthogonal sections at *s*.

Shear variables and bending strains are coupled

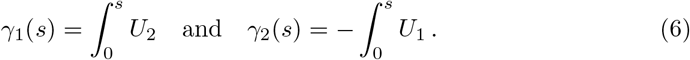

The above formulas, which result from the constrained kinematics of the axonemal structure (as explained further in Appendix A), underlie the essential mechanism of axonemal motility: collective sliding of MTs generates bending of the whole Ax. We point out here that there is no coupling between the shear variables *γ*_1_, *γ*_2_ and the twist *U*_3_, a fact that will have consequences in the remainder.

The special axonemal deformation in Figure 4 shows the case in which *U*_1_(*s*) = 0 and *U*_2_(*s*) = *K*, so the Ax is bent into a circular arc of radius 1/*K*. While *γ*_2_(*s*) = 0, the shear variable *γ*_1_(*s*) = *K*_*s*_ increases linearly with *s*. Material and orthogonal sections coincide at the base (the basal body impose no shear at *s* = 0) and the angle between them grows as we move along the centerline towards the distal end of the Ax. In order for the Ax to bend, MTs from one side of the Ax must be driven driven towards the distal end while the others must be driven toward the proximal end.

We remark here that (4) defines the most general active internal energy generated by molecular motors, and we do not assume at this stage any specific (spatial) organization of dynein forces. We will introduce specific shear forces later in the Results Section.

The PFR is modeled as an elastic cylinder with circular cross sections of radius *ρ*^*p*^ and rest length *L*. We assume that the PFR can stretch and shear. The total internal energy of the PFR is given by

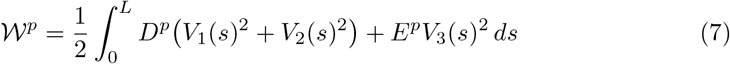

where *V*_1_ and *V*_2_ are the shear strains, *V*_3_ is the stretch, *D*^*p*^ and *E*^*p*^ are the shear and stretching moduli, respectively. We are neglecting here the PFR’s bending and twisting resistance. Classical estimations on homogeneous elastic rods, see e.g. [13], show that bending and twist moduli scale with the forth power of the cross section radius, whereas shear and stretching moduli scale with the second power and hence they are dominant for small radii. We assume that dynein forces are strong enough to induce shear in the PFR, thus PFR’s bending and twist contributions to the energy of the flagellum become negligible. We are also neglecting Poisson effects by treating the PFR cross sections as rigid.

The PFR shear strains and stretch are defined as follows. The cross sections centers of the PFR lie on the curve **r**^*d*^, and their orientations are given by the orthonormal frame **g**_*i*_(*s*), with *i* = 1, 2, 3. The unit vectors **g**_1_(*s*) and **g**_2_(*s*) determine the cross section plane centered at **r**^*p*^(*s*), while the unit vector **g**_3_(*s*) is orthogonal to it. The curve **r**^*p*^ is not parametrized by arc length and **g**_3_ is not in general aligned with the tangent to **r**^*p*^. Shear strains and stretch are given by the formulas

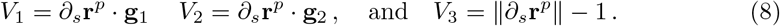

The shear strains thus depend on the orientation of the cross sections with respect to the centerline (tangent), while the stretch measures the elongation of the centerline.

The PFR-Ax attachment couples the kinematics of the two substructures, see Figure 4. In the remainder we formalize the attachment constraint and we show how the PFR’s shear strains and stretch (8), and thus the flagellar energy (1), are completely determined by the Ax kinematic variables.

For each *s*, the PFR cross section centered at **r**^*p*^(*s*) is in contact with the Ax surface at the point **C**(*s, ϕ*^*p*^) for for a fixed angle coordinate *ϕ*^*p*^, see Figure 1. The PFR centerline is given by

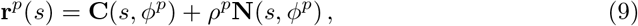

where **N**(*s, ϕ*^*p*^) ≈ **d**_1_(*s*) cos *ϕ*^*p*^ + **d**_2_(*s*) sin *ϕ*^*p*^ is the outer unit normal to the axonemal surface at **C**(*s, ϕ*^*p*^). The normal vector **N**(*s, ϕ*^*p*^) lies on the plane of the PFR cross section centered at **r**^*p*^(*s*). Indeed, we have **g**_1_(*s*) = **N**(*s, ϕ*^*p*^) for the first unit vector of the PFR orthonormal frame. Only one more degree of freedom remains, namely **g**_2_(*s*), which must be orthogonal to **N**(*s, ϕ*^*p*^), to fully characterize the orientations of the PFR cross sections. Here is where the bonding links attachments are introduced in the model. The bonding links of the PFR cross section centered at **r**^*p*^(*s*) are attached to three adjacent MTs at the same MTs’ arc length *s*. The individual attachments are therefore located on the material section of the Ax at *s*. Given this, **g**_2_(*s*) is imposed to be parallel to *∂*_*ϕ*_**C**(*s, ϕ*^*p*^), the tangent vector to the material section of the Ax at the point of contact **C**(*s, ϕ*^*p*^), see Figure 4. This condition critically couples MTs’ shear to the orientations of the PFR cross sections, as further demonstrated below.

To summarize, we have the following formulas for the PFR orthonormal frame vectors

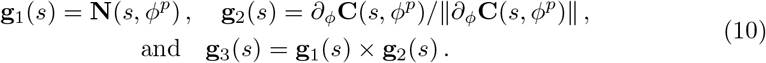

By replacing the expression (9)-(10) in (8), we obtain formulas for the shear strains and stretch of the PFR in terms of the Ax kinematic parameters. We have that the shear strain *V*_1_ and the stretch *V*_3_ are of order *ρ*^*p*^ ~ *ρ*^*a*^ (see Appendix A for detailed calculations). Since *ρ*^*p*^ is small compared to the length scale *L* of both PFR and Ax we can neglect these quantities. The only non-negligible contribution to the PFR energy is thus given by the shear strain *V*_2_. After linearization, we have *V*_2_ ≈ −sin *ϕ*^*p*^*γ*_1_ + cos *ϕ*^*p*^*γ*_2_. The PFR energy in terms of Ax kinematic parameters is then given by

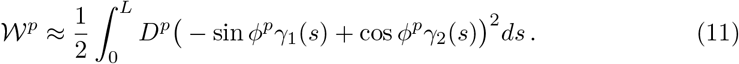

The shear of axonemal MTs determines the orientation of the PFR cross sections. In Figure 5 (middle pictures) we show an example of this kinematic interplay. The Ax is again bent in an arc of a circle on the plane **d**_1_ − **d**_3_, with *U*_1_(*s*) = 0, *U*_2_(*s*) = *K*, *γ*_1_(*s*) = *K*_*s*_, and *γ*_2_(*s*) = 0. PFR and Ax centerlines run parallel to each other, indeed from (9) we have that *∂*_*s*_**r**^*a*^ ≈ *∂*_*s*_**r**^*p*^ for every deformation. The linking bonds impose a rotation of the cross sections of the PFR as we progress from the proximal to the distal end of the flagellum, generating shear strain *V*_2_(*s*) = −sin *ϕ*^*p*^*γ*_1_(*s*) = −sin *ϕ*^*p*^*K*_*s*_ on the PFR. This mechanical interplay leads to non-planarity of the euglenid flagellar beat. This mechanism is controlled by the offset between the PFR-Ax joining line and the local spontaneous bending plane of the Ax, as further discussed in the Results Section.

**Figure 5.**
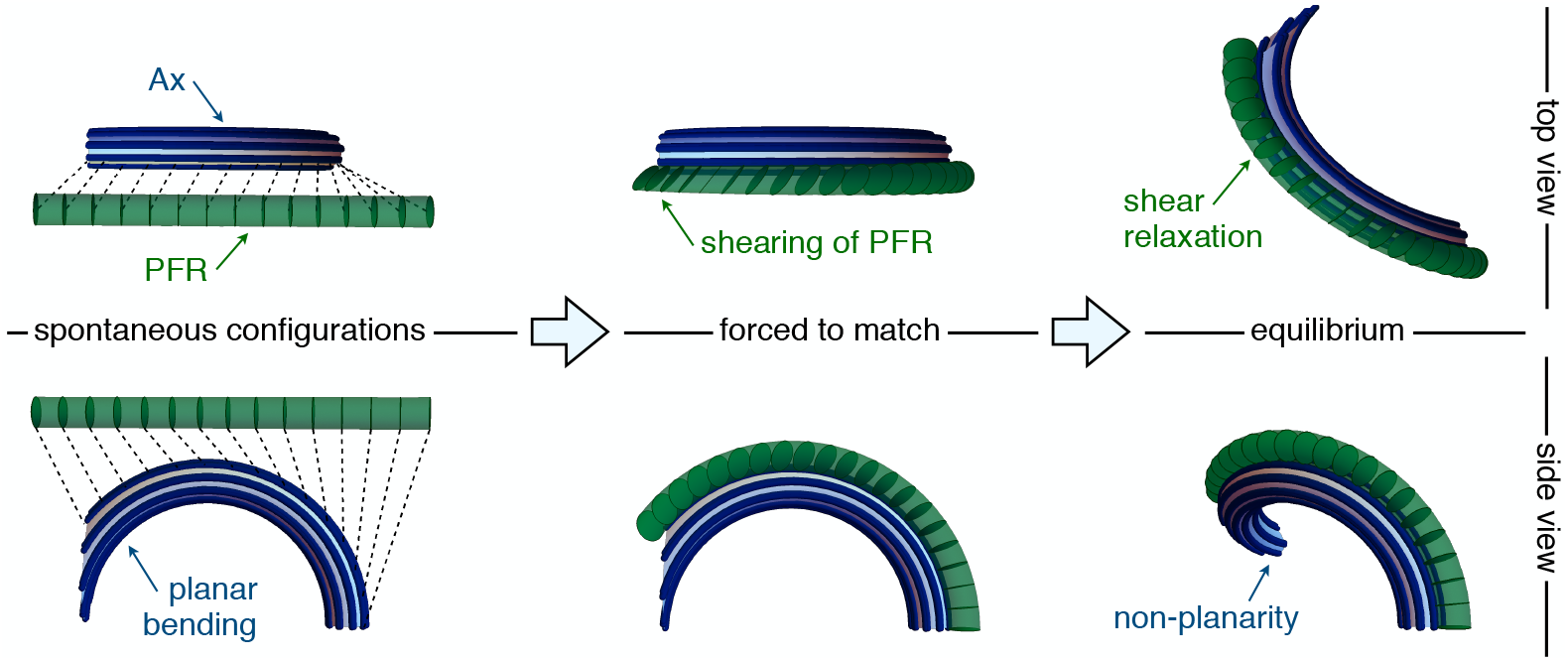
Flagellar non-planarity arising from structural incompatibility. The Ax-PFR mechanical interplay is explained in a three-steps argument (left to right). Consider first the two separated structures in their spontaneous configurations (left). The Ax is bent into a planar arc while the PFR is straight. Then, the PFR is forced to match to the Ax, while the latter is kept in its spontaneous configuration (middle). The attachment constraint induces shear strains in the PFR. The composite system cannot then be in mechanical equilibrium without external forcing. When the composite system is released (right), it reaches equilibrium by the relaxation of the PFR shear, which induce additional distortion of the Ax. At equilibrium, an optimal energy compromise is reached, which is characterized by an emergent non-planarity.

### Equilibria

Under generic (steady) dynein actuation, i.e. given *H*_1_ and *H*_2_ (not time-dependent), and in the absence of external forces, the flagellum deforms to its equilibrium configuration 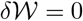. Bending strains and twist at equilibrium solve the equations

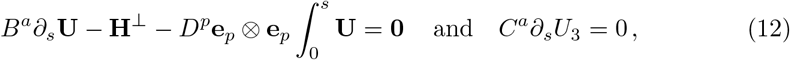

where **U** = (*U*_1_, *U*_2_) is the bending vector, **e**^*p*^ = (cos *ϕ*^*p*^, sin *ϕ*^*p*^), and **H**^⊥^ = (−*H*_2_, *H*_1_). We use the symbol **a** ⊗ **b** to denote the matrix with components (**a** ⊗ **b**)_*ij*_ = *a*_*i*_*b*_*j*_. The field equations (12) are complemented by the boundary conditions

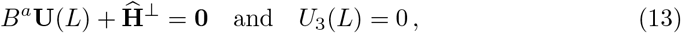

where 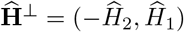. Equations (12)-(13) can be interpreted as the torque balance equations of the Ax. The derivative of the (elastic) bending moment and the internal shear stresses balance the torque per unit length exerted by the PFR on the Ax, which is given by the *D*^*p*^-dependent term appearing in the first equation. The torque depends on the integral of the bending vector, making the balance equations non-standard (integrodifferental instead of differential). This dependency is due to the fact that the torque arises from the shear deformations of the PFR, which are induced by the shear of axonemal MTs, which is, in turn, related to axonemal bending strains via the integral relations (6). The torque exerted by the PFR on the Ax is sensitive to the direction given by the unit vector **e**_*p*_, hence it depends on the angle *ϕ*^*p*^ between the Ax-PFR joining line and the unit vector **d**_1_.

### Hydrodynamics

We derive here the dynamic equations for a flagellum beating in a viscous fluid. We consider the extended functional

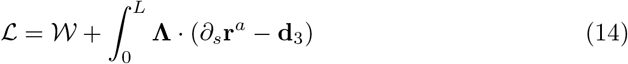

where **Λ** is the Lagrange multiplier vector enforcing the constraint *∂*_*s*_**r**^*a*^ = **d**_3_. We treat the fluid-flagellum interaction in the local drag approximation of Resistive Force Theory, see e.g. [35]. In this approximation, viscous forces and torques depend locally on the translational and rotational velocity of the flagellum, represented here for simplicity by the translational and rotational velocity of the Ax. The external viscous forces **F** and torques **G** (per unit length) acting on the flagellum are given by

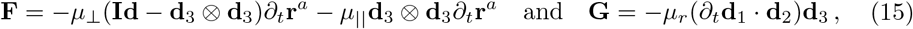

where *μ*_⊥_, *μ*_‖_, and *μ*_*r*_ are the normal, parallel, and rotational drag coefficient (respectively), and **Id** is the identity tensor. The principle of virtual work imposes

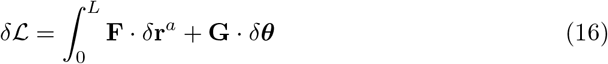

for every variation *δ***r**^*a*^ and *δθ* = *δθ*_1_**d**_1_ + *δθ*_2_**d**_2_ + *δθ*_3_**d**_3_, where *δθ*_1_ = (*δ***d**_2_ · **d**_3_), *δθ*_2_ = (*δ***d**_3_ · **d**_1_), and *δθ*_3_ = (*δ***d**_1_ · **d**_2_). Linearizing the force balance equations derived from (16) we obtain the following equations for bending strains and twist

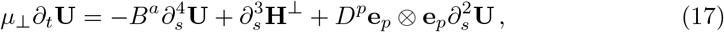

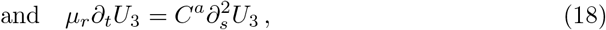

which are decoupled from the extra unknown **Λ**. Equations (17) and (18) are complemented by the boundary conditions

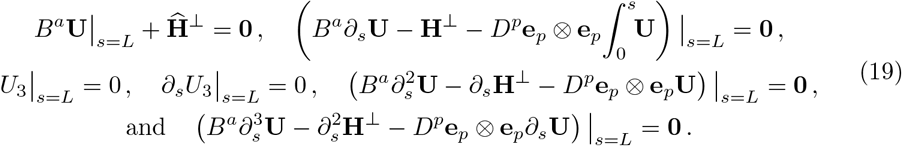

The details of the derivation of (17)-(19) are provided in Appendix B.

Once we solve for *U*_1_, *U*_2_, and *U*_3_ either the equilibrium equations (12)-(13) or the dynamic equations (17)-(19), the shape of the flagellum can be recovered. In particular, we obtain the orthonormal frame **d**_*i*_ with *i* = 1, 2, 3 by solving while the centerline of the Ax is recovered by integrating *∂*_*s*_**r**^*a*^ = **d**_3_.

## Results

We analyze the geometry of the centerline **r**^*a*^ which, due to the slenderness of the flagellar structure, is a close proxy for the shape of the flagellum.

In general, the shape of a curve is determined by its curvature *κ* and torsion *τ*. Since **r**^*a*^ is parametrized by arc length, the two quantities are given by the formulas *∂*_*s*_**t** = *κ***n** and *∂*_*s*_**b** = −*τ***n**, where **t** = *∂*_*s*_**r**^*a*^, **n** = *∂*_*s*_**t**/|*∂*_*s*_**t**|, and **b** = **t × n** are the tangent, normal, and binormal vector to the curve **r**^*a*^, respectively. Given *κ* and *τ*, **r**^*a*^ is uniquely determined up to rigid motions.

From the previous definitions and from (3) we obtain the relations between curvature, torsion, bending, and twist. In compact form these relations are given by

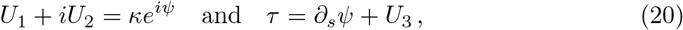

which hold for **U** ≠ 0. In (20) we introduced the angle *ψ* that the bending vector **U** = (*U*_1_, *U*_2_) forms with the line *U*_2_ = 0, see Figure 6. Now, at equilibrium (12) we have

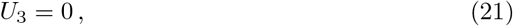

under any dynein actuation. In other words, axonemal deformations are twistless. This is, fundamentally, a consequence of the fact that shear of axonemal MTs and twist are uncoupled (6). The torsion of the centerline **r**^*a*^ is our main focus, since we are interested in emergent non-planarity. Combining (20) and (21) we have that torsion can arise only from the rotation rate *∂*_*s*_*ψ* of the bending vector **U** along the length of the flagellum. This last observation will be important in the following.

**Figure 6.**
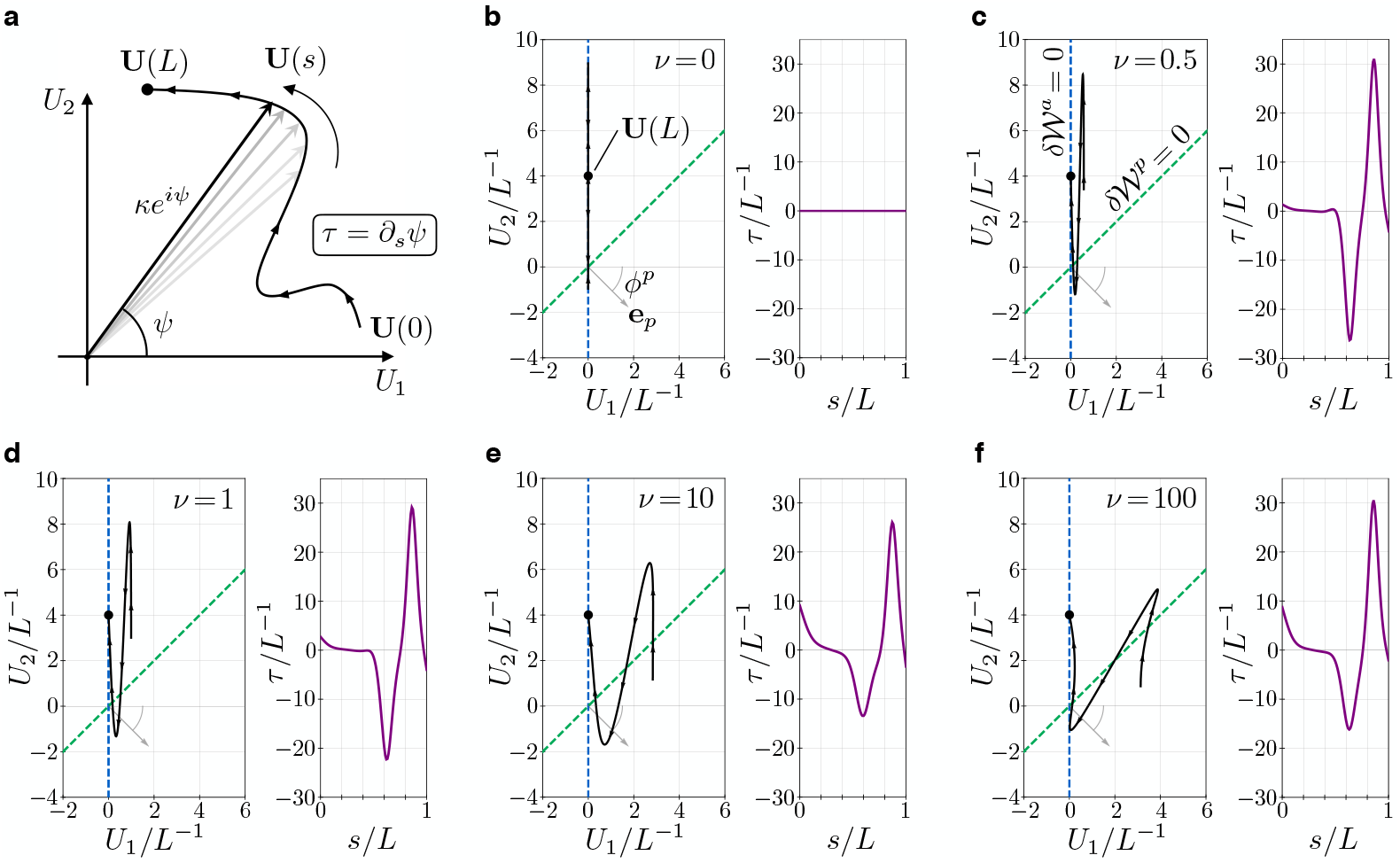
a) The bending vector **U**(*s*) = (*U*_1_(*s*), *U*_2_(*s*)) traces a curve on the plane of the bending parameters *U*_1_ and *U*_2_. The norm of the bending vector determines the curvature *κ*(*s*) = |**U**(*s*)| of the flagellum. The rate of change of the angle *ψ*(*s*) determines the torsion *τ* = *∂*_*s*_*ψ*. b-f) Bending vectors of flagellar equilibrium configurations under the same (steady) dynein actuation, but different values of the material parameter *ν* = *D*^*p*^/(*B*^*a*^*L*^−2^). Equilibria are minimizer of the energy 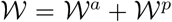. For small values of *ν* the Ax component of the energy 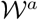 dominates. In this case **U** is close to the target bending vector 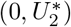 where 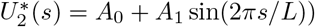. For large values of *ν* the PFR component of the energy 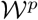 dominates, and equilibria are dragged closer to the line orthogonal to the vector **e**_*p*_ (dashed green). The bending vector undergoes rotations which result in torsional peaks of alternating sing.

### Dynein actuation induced by sliding inhibition

Under the assumptions (21) and (23), the flagellar energy (1) can be rewritten as

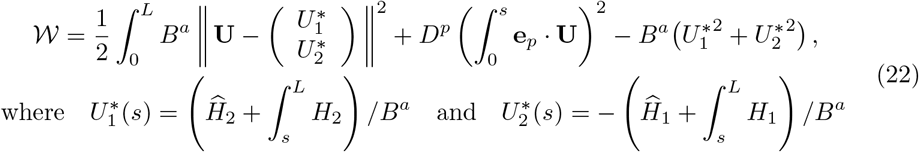

are the target bending strains generated by the dynein forces. The use of this terminology is clear from (22). The effect of dynein actuation at equilibrium (when the energy is minimized) is to bring the bending strains *U*_1_ and *U*_2_ as close as possible to 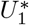 and 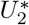, respectively. The emerging bending strains and the target bending strains might not match due to the interference by the PFR component of the energy (*D*^*p*^ ≠ 0). From the formulas for the target bending strains in (22) we can infer the importance of the concentrated shear forces 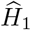 and 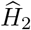. Without these forces, admissible spontaneous configurations of the Ax would be ruled out. If the concentrated shear forces are null, for example, the Ax cannot spontaneously bend into a circular arc. Indeed, for a circular arc of radius 1/*K* on the plane **d**_1_-**d**_3_ we must have 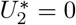 and 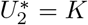. In this case, from (22) we have that 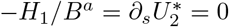, which implies *H*_1_ = 0 and 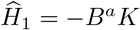, so the concentrated forces must be non zero.

As per our main hypothesis, we suppose that the sliding inhibition exerted by the bonding links let dyneins organize so that the Ax locally bends spontaneously on the plane **d**_1_(*s*)-**d**_3_(*s*), as shown in Figure 1. This is equivalent to require that 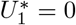, which leads to the following condition on the shear forces

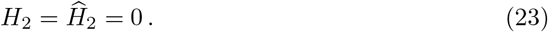

### Emergence of non-planarity

We consider here the equilibrium equations (12) under the hypothesis (23). We look at the equilibrium configurations for every possible value of the angle *ϕ*^*p*^ between the Ax-PFR joining line and the spontaneous bending plane of the Ax, even though the value of actual interest for Euglena is *ϕ*^*p*^ ≈ −2*π*/9. We can prove analytically the following statement: *if the Ax-PFR joining line is neither parallel nor orthogonal to the spontaneous bending plane of the Ax, then the emergent flagellar shapes are non-planar*.

Indeed, suppose *H*_1_ ≠ 0. From (20) and (21) it follows that the shape of the flagellum is planar *τ* = 0 if (and only if) the angle *ψ* of the bending vector **U**(*s*) = (*U*_1_(*s*), *U*_2_(*s*)) is constant. The bending vector must therefore be confined on a line for every *s*. In this case there must be two constants *c*_1_ and *c*_2_ such that *U*_1_(*s*) = *c*_1_*U* (*s*) and *U*_2_(*s*) = *c*_2_*U* (*s*) for some scalar function *U*. Now, if a planar **U** is a solution of (12) then we must have

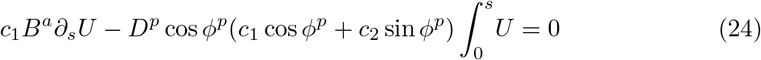

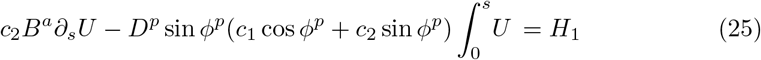

with *c*_1_*U*(*L*) = 0 and 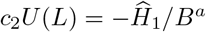. If *ϕ*^*p*^ ≠ 0, *π*/2, *π*, 3*π*/2 the system of equations (24)-(25) admits no solution. Indeed, suppose first that 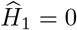. Since *H*_1_ ≠ 0 we must have (*c*_1_, *c*_2_) ≠ (0, 0). However, in this case, (24) admits the unique solution *U* = 0, which is incompatible with (25). If 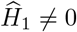 then the boundary conditions impose *c*_1_ = 0, but in this case (24) has again *U* = 0 as a unique solution, which is incompatible with both the boundary conditions and with (25). Our statement is thus proved.

For *ϕ*^*p*^ ≈ −2*π*/9, the characteristic value for Euglena, the non-planarity of flagellar shapes is not just possible. It is the only possible outcome under any non-trivial dynein actuation.

### Structural incompatibility and torsion with alternating sign

Alongside the previous analysis there is a less technical way to infer the emergence of non-planarity from our model. We look here more closely to the flagellum energy, and we think in terms of structural incompatibility between Ax and PFR, seen as antagonistic elements of the flagellum assembly, see Figure 5.

Under the assumptions (21) and (23), the flagellar energy is given by

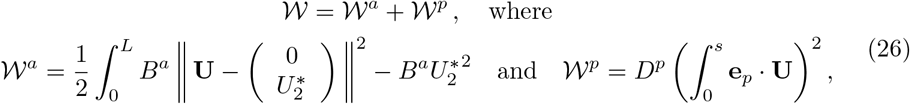

with 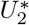 as in (22). The energy has two components, 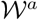 that depends on the Ax bending modulus *B*^*a*^, and 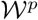 that depends on the PFR shear modulus *D*^*p*^. We can vary these material parameters and explore what the resulting minima of 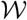, i.e. the equilibrium configurations (12)-(13), must look like. We consider the nondimensional parameter *ν* = *D*^*p*^/(*B*^*a*^*L*^−2^). When *ν* ≪ 1 the Ax component 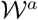 of the energy dominates. In this case, at equilibrium, the bending vector has to be close to the target bending vector 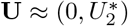. In particular, then, **U**(*s*) will be confined near the line *U*_1_ = 0 for every *s*. In the case *ν* ≫ 1 the PFR component 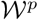 dominates, and the energy is minimized when **U**(*s*) lies close to the line generated by the vector 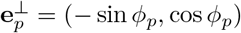. Clearly, if the latter line is different from *U*_1_ = 0, the two extreme regimes *ν* ≪ 1 and *ν* ≫ 1, each of which favours one of the two individual components, aim at two different equilibrium configurations. In other words, Ax and PFR are structurally incompatible.

When neither of the two energy components dominates, the emergence of non-planarity can be intuitively predicted with the following reasoning. In the intermediate case *ν* ~ 1 we expect the equilibrium configurations to be a compromise among the two extreme cases, with the bending vector **U**(*s*) being “spread out” in the region between the two extreme equilibrium lines. The spreading of the bending vector is aided by the concentrated shear force at the tip, which imposes 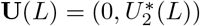 irrespectively of the PFR stiffness. The bending vector is then “pinned” at *s* = *L* on the *U*_1_ = 0 line while it gets dragged toward the line generated by 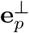 for large values of *ν*. Hence the spreading. The bending vector will then span an area and, consequently, undergo rotations. Since torsion is determined by the rotation rate of the bending vector *τ* = *∂*_*s*_*ψ*, the resulting flagellar shapes will be non-planar.

Figure 6 illustrates a critical example in which the previous intuitive reasoning effectively plays out. We consider a target bending of the kind 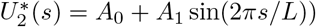, a fair idealization of the asymmetric shapes of a Chlamydomonas-like flagellar beat [10]. We take *ϕ*^*p*^ = −*π*/4 (larger than the Euglena value, to obtain clearer graphs). For *ν* = 0 the bending vector lies inside the *U*_1_ = 0 line, and its amplitude oscillates. For positive values of *ν*, when the PFR stiffness is “turned on”, the oscillating bending vector is extruded from the *U*_1_ = 0 line. For large values of *ν* it gets closer and closer to the line generated by 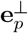. The bending vector spans an area and, following the oscillations, it rotates clock-wise and anti-clock-wise generating an alternation in the torsion sign. This is the geometric signature of the spinning lasso.

### Hydrodynamic simulations and comparison with observations

Our model is able to predict the torsional characteristic of the euglenid flagellum in the static case, and in absence of external forces. We test here the model in the more realistic setting of time-dependent dynein actuation in the presence of hydrodynamic interactions.

We first observe that, as in the static case, the dynamic equations for **U** and *U*_3_ are decoupled (17)-(18), and that dynein forces do not affect twist. We have then twistless kinematics under any actuation also in the dynamic case, at least after a time transient. We can simply assume (21) for all times, so the torsion of **r**^*a*^ is still completely determined by the bending vector.

We consider a dynein actuation that generates Chlamydomonas-like shapes in a flagellum with no extra-axonemal structures. The shear forces *H*_1_ and 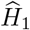 employed in our simulation are shown in Figure 7. The same figure also shows the emergent bending strains of a PFR-free flagellum actuated by said forces, beating in a viscous fluid. The dynamic equations for this system are simply (17)-(18) with *D*^*p*^ = 0. The resulting bending strains, which generate a planar beat, resembles the experimentally observed Chlamydomonas flagellar curvatures reported in [23].

**Figure 7.**
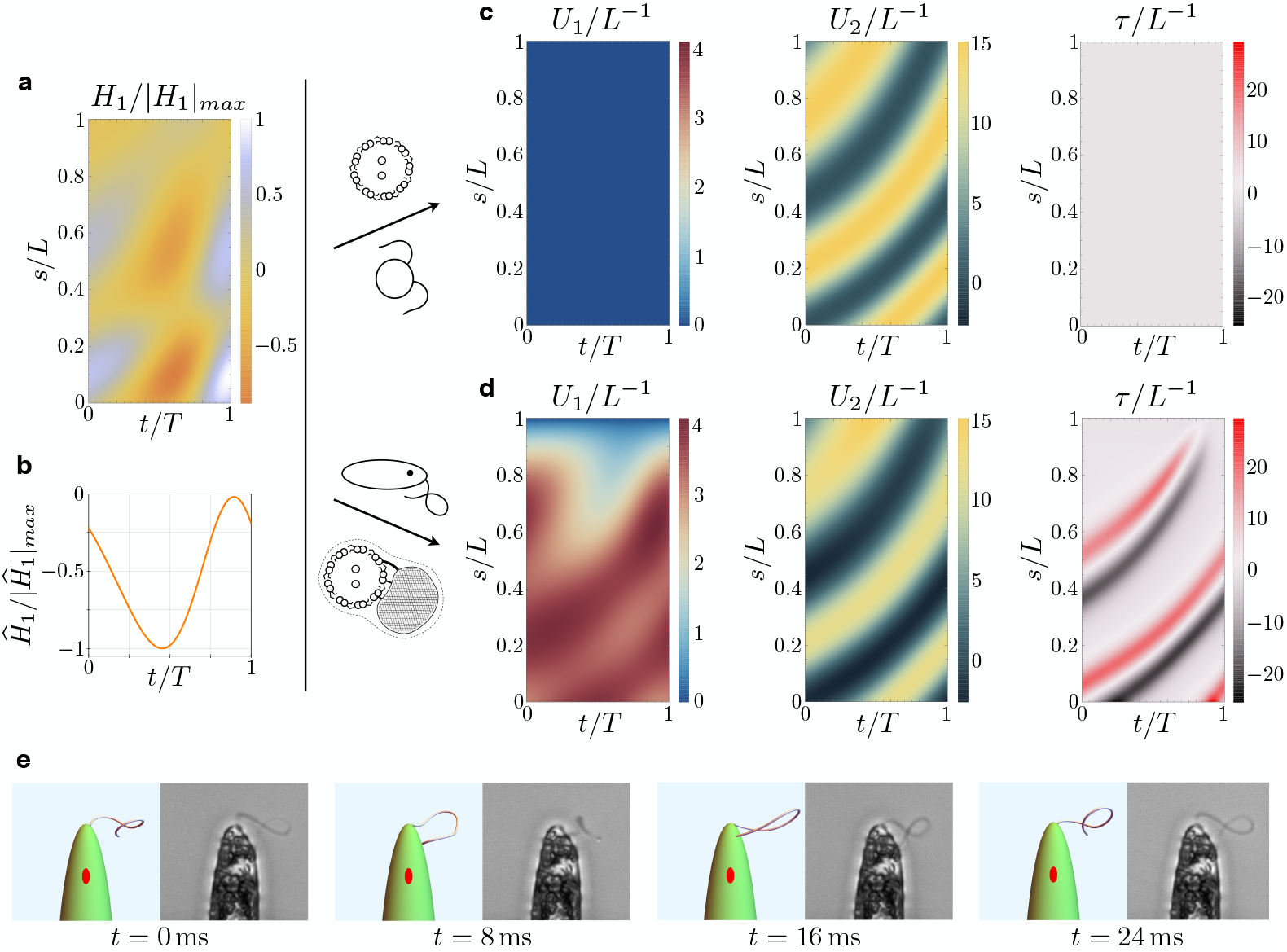
a-b) Dynein shear forces. c) Resulting bending strains and torsion for an Ax actuated by the force pattern (a-b), beating in a viscous fluid, and free of extra-axonemal structures. The beat is planar (Chlamydomonas-like). d) Resulting bending strains and torsion for an euglenid flagellum (composite structure Ax+PFR) actuated by (a-b) and beating in a viscous fluid. The Ax-PFR interaction generates torsional peaks with alternate sign travelling from the proximal to the distal end of the flagellum. e) Resulting shapes for the euglenid flagellum at different instants within a beat, and comparison with experimental observations.

Finally, Figure 7 presents the emergent bending strains of the beating euglenid (PFR-bearing) flagellum, together with the corresponding flagellar torsion. The spinning lasso torsional signature is clearly present. Indeed, the fluid-structure interaction does not disrupt the Ax-PFR structural incompatibility, which still generates non-planar shapes with travelling waves of torsional peaks with alternating sign and the typical looping-curve outlines, cfr. Figure 2. All the details on the methods and parameters employed in the simulations are given in Appendix C and Appendix D.

## Discussion and Outlook

We have shown how the origin of the peculiar shapes of the euglenid flagellum can be explained by the mechanical interplay of two antagonistic flagellar components, the Ax and the PFR. Our conclusions are based mainly on the hypothesis that sliding inhibition by the PFR organizes dynein activity, and localizes the spontaneous bending plane of the Ax as the one that passes from the Ax center through the MTs bonded to the PFR. This is in agreement with the current understanding of the mechanism that generates beat planarity in other PFR-bearing flagellar systems. Non-planarity in Euglena can arise because of a marked asymmetry in the Ax-bonding links-PFR complex in the Euglenid flagellum, which is not found in kinetoplastic organisms such as Leishmania [11] or Trypanosoma [22].

In the absence of a precise knowledge of the dynein actuation pattern we tested our mechanical model under shear forces that would, in the absence of extra-axonemal structures, realize a beat similar to those found in model systems like Chlamydomonas. We appreciate that the emergent distortion of the Ax, generated by the Ax-PFR interplay, could in principle lead to different actuation patterns, consistently with the hypothesis of dynein actuation via mechanical feedback. This question will require further studies.

Along with the mechanism that let the euglenid flagellar shapes emerge, it is worth considering how this characteristic flagellar beat is integrated in the overall behaviour of the organism. As shown in [25], the spinning lasso beat produces the typical roto-translational trajectories of swimming euglenas. Cell body rotation is in turn associated with phototaxis. Indeed, rotation allows cells to “scan” the environment, and veer to the light source direction when stimulated (or escape in the opposite direction, when the signal is too strong). Here, the key biochemical mechanism could be the one often found in nature, by which periodic signals generated by lighting and shading associated with body rotations are used for navigation, in the sense that the existence of periodicity implies a lack of proper alignment [12]. It is known that the PFR is directly connected with the light-sensing apparatus [24], and might even be active [21]. Further study on euglenid flagellar motility and phototaxis could lead to a more comprehensive understanding on the role of the PFR.

## Materials and Methods

Strain SAG 1224-5/27 of *Euglena gracilis* obtained from the SAG Culture Collection of Algae at the University of Göttingen was maintained axenic in liquid culture medium Eg. Cultures were transferred weekly. Cells were kept in incubator at 15° C at a light:dark cycle of 12:12 h under a cold white LED illumination with an irradiance of about 50 *μ*mol · m^−2^ · s^−1^.

An Olympus IX 81 inverted microscope with motorized stage was employed in all the experiments. Experiments were performed at the Sensing and Moving Bioinspired Artifacts Laboratory of SISSA. The microscope was equipped with a LCAch 20X Phc objective (NA 0.40) for the imaging of cells trapped at the tip of a glass capillary using transmitted brightfield illumination. The intermediate magnification changer (1.6 X) of the microscope was exploited to achieve higher magnification. Micrographs were recorded at a frame rate of 1, 000 fps with a Photron FASTCAM Mini UX100 high-speed digital camera.

Tapered capillaries of circular cross section were obtained from borosilicate glass tubes by employing a micropipette puller and subsequently fire polished. At each trial observation a glass capillary was filled with a diluted solution of cells and fixed to the microscope stage by means of a custom made, 3d-printed holder. The holder allowed for keeping the capillary in place and rotating it about its axis, so as to image a cell specimen from distinct viewpoints. Cells were immobilized at the tip of the capillary by applying a gentle suction pressure via a syringe connected to the capillary by plastic tubing.

## Acknowledgments

We thank A. Beran for his assistance with *E. gracilis* samples. This study was supported by the European Research Council through the ERC Advanced Grant 340685-MicroMotility.

## A Model details

The Ax consists of a bundle of inextensible filaments of length *L* (MTs) lying on a cylindrical surface of radius *ρ*^*a*^. For simplicity, the model ignores the mechanical effects of radial spokes and the central pair. The axonemal surface is parametrized by the generalized cylindrical coordinates *z* and *ϕ* via the map

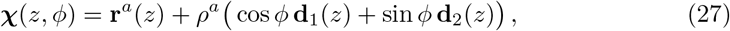

with **r**^*a*^, **d**_1_, and **d**_2_ defined as in the main text. Following [14], we suppose that the axonemal constraints confine MTs on the Ax surface at a fixed angular distance Δ*ϕ* = 2*π*/9 between each other. More formally, for *j* = 1, . . . , 9, we define the centerline **r**^*j*^ of the *j*-th MT as **r**^*j*^(*s*) = **C**(*s, ϕ*_*j*_), where

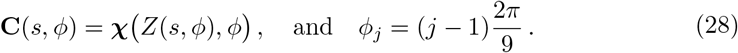

The function *Z*(*s, ϕ*) in (28) is defined (implicitly) via the equality

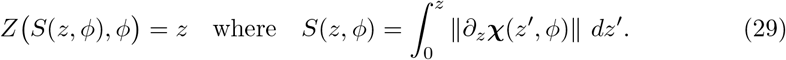

From the definitions above follows ‖*∂*_*s*_**r**^*j*^‖ = 1, so MTs are indeed inextensible and *s* is their arc length. Moreover, the Taylor expansion of **C** at the first order in *ρ*^*a*^ gives the approximated formula (5), with *γ*_1_ and *γ*_2_ given by (6).

We associate to the *j*-th MT an orthonormal frame along **r**^*j*^ given by the unit vectors

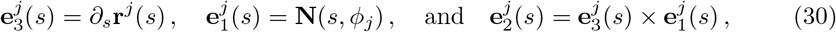

where **N** = cos *ϕ* **d**_1_(*Z*) + sin *ϕ* **d**_2_(*Z*) is the (outer) unit normal to the cylindrical surface. The unit vectors (30) determine MTs’ cross section orientations. The unit vectors 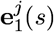 and 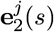 lie on the cross-section centered at **r**^*j*^(*s*), while 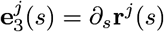 is orthogonal to it. The (passive) elastic energy of the Ax is given by the sum of the MTs’ elastic energies

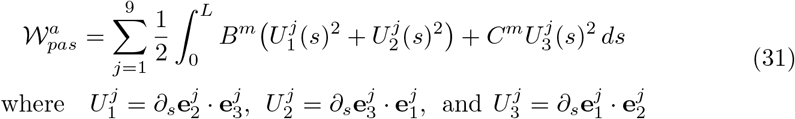

are the strains associated to the *j*-th MT, while *B*^*m*^ and *C*^*m*^ are the MTs’ bending and twisting moduli (respectively). At the leading order approximation in *ρ*^*a*^ we have

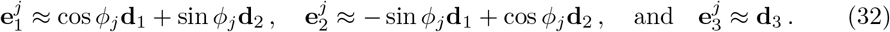

From (32) and (3) follows that 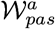, as defined in (31), can indeed be approximated by the right hand side of (2), with *B*^*a*^ = 9*B*^*m*^ and *C*^*a*^ = 9*C*^*m*^.

We consider now MT sliding. Fixed a point **r**^*j*−1^(*s*) on the (*j* − 1)-th MT’s centerline we look for the nearest point to **r**^*j*−1^(*s*) on the centerline **r**^*j*^ of the *j*-th MT. Such a point **r**^*j*^(*s*∗) (we can think of it as a projection) lies at some arc length *s*∗, which depends on *s*. We write *s*∗ = Π_*j*_(*s*). We then define the sliding *σ*_*j*_(*s*) as the difference of the two arc lengths *s* and Π_*j*_(*s*). More formally,

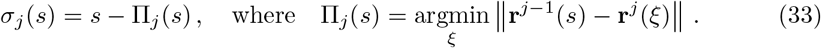

Figure A1 illustrates the geometric idea of definition (33).

**Figure A1.**
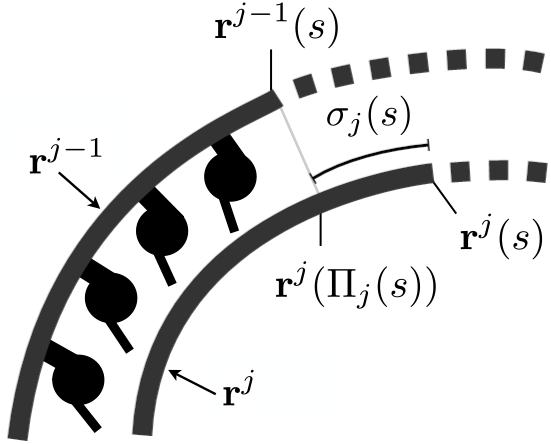
Sketch of two MTs’ centerlines during deformation. The sliding *σ*_*j*_(*s*) is defined as the difference between the arc lengths *s* and Π_*j*_(*s*). The latter is the arc length corresponding to the projection of **r**^*j*−1^(*s*) on the curve **r**^*j*^. We have positive sliding when dyneins push the *j*-th MT towards the distal end of the flagellum and the (*j* − 1)-th MT toward the proximal end.

The active internal energy of the Ax is defined as minus the total mechanical work of the dyneins

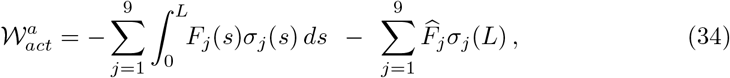

where *F*_*j*_(*s*) are the sliding forces on the *j*-th MT exerted by the dyneins on the (*j* − 1)-th MT, and 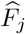 are the singular sliding forces (on the *j*-th MT exerted by the dyneins on the (*j* − 1)-th MT) concentrated at the distal end of the Ax. Taylor expanding (33) in *ρ*^*a*^ we have, at the leading order,

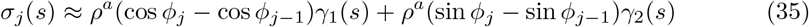

with *γ*_1_ and *γ*_2_ given by (6). From (35) we have that 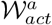, as defined in (34), is approximated by the right hand side of (4), with

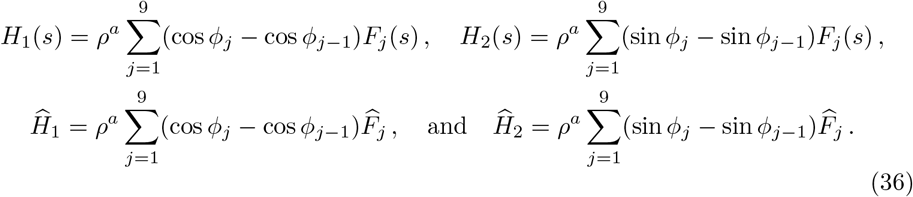

In the remainder we give some details of the derivation of (11) from (7). Expanding (10) at the leading order in *ρ*^*p*^ gives

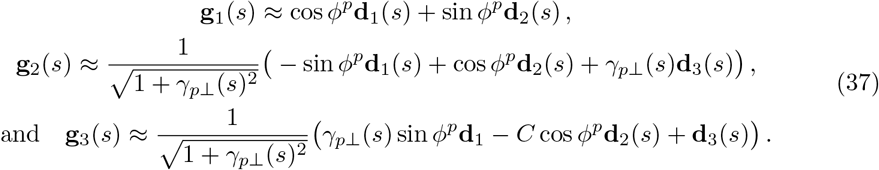

with *γ*_*p*⊥_(*s*) = − sin *ϕ*^*p*^*γ*_1_(*s*) + cos *ϕ*^*p*^*γ*_2_(*s*). From (8) and (37), leading order calculations *V*_1_, *V*_3_ ~ *ρ*^*p*^, whereas 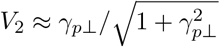. Linearizing in *γ*_*p*⊥_ we have *V*_2_ ≈ *γ*_*p*_, from which follows (11).

## B Dynamical equations

To derive equations (17)-(19) it is convenient to introduce the quantities *M*_1_, *M*_2_, and *M*_3_ defined via the following variational equality

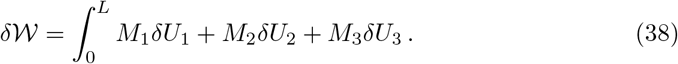

These quantities can be interpreted as the local components of the flagellar moment

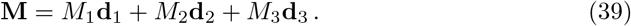

A direct calculation gives

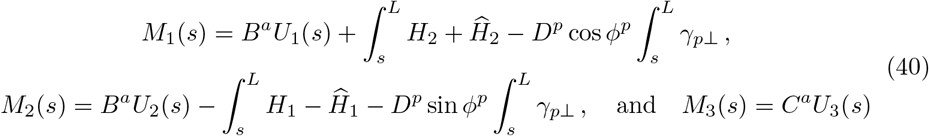

where *γ*_*p*⊥_(*s*) = − sin *ϕ*^*p*^*γ*_1_(*s*) + cos *ϕ*^*p*^*γ*_2_(*s*), as in the previous section. We then write the variations *δU*_*i*_ in terms of *δθ*_*i*_ (defined in the main text) obtaining

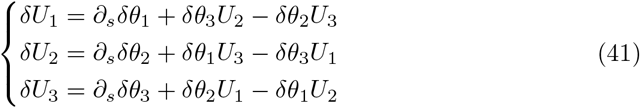

Combining (38), (39) and (41) we have

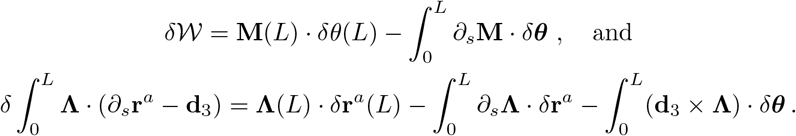

In the calculations above we took variations with *δ***r**^*a*^(0) = *δθ*(0) = **0**, since we consider a flagellum with a clamped end at *s* = 0. The principle of virtual work (16) gives us then the following force and torque balance equations

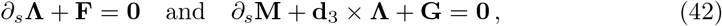

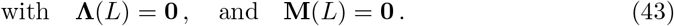

Equations (17)-(19) are derived from (42) and (43), after some extra formal manipulations that we explain in the reminder.

We first introduce the local angular velocities

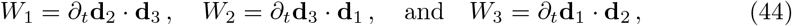

which are related to the strains via the following compatibility equations

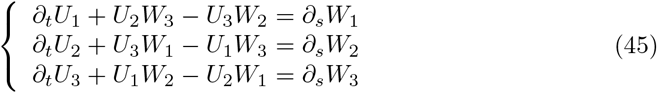

Equations (45) follow from the identities *∂*_*s*_*∂*_*t*_**d**_*i*_ = *∂*_*t*_*∂*_*s*_**d**_*i*_, with *i* = 1, 2, 3. We then rewrite the external forces and torques (15) in compact form as

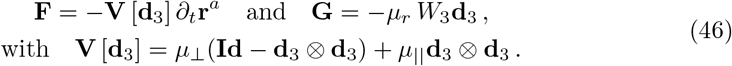

Scalar multiplying the torque balance equation by **d**_1_ and **d**_2_ we obtain the expressions for the first two local components

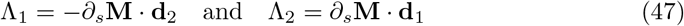

of the Lagrange multiplier vector **Λ** = Λ_1_**d**_1_ + Λ_2_**d**_2_ + Λ_3_**d**_3_. Scalar multiplying the torque balance equation by **d**_3_ gives

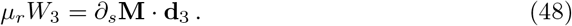

We rewrite the force balance equation as *∂*_*t*_**r**^*a*^ = **V**[**d**_3_]^−1^ *∂*_*s*_**Λ** and, after differentiating both sides with respect to *s* and then scalar multiplying by **d**_1_, **d**_2_, and **d**_3_, we have

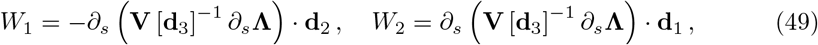

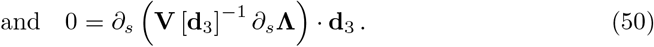

From (40), (45), and (48) we obtain (18) by first differentiating with respect to *s* both sides of (48), and then by linearizing the resulting equation. Similarly, exploiting also equations (47) this time, we differentiate with respect to *s* and then linearize (49) to obtain (17). Boundary conditions (19) follow from (43) and

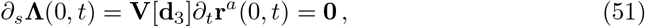

which follows from the fixed end condition **r**^*a*^(0, *t*) = **0**.

Given *U*_1_, *U*_2_, and *U*_3_ the orthonormal frame **d**_1_, **d**_2_, and **d**_3_ can be recovered by integrating the system of equations

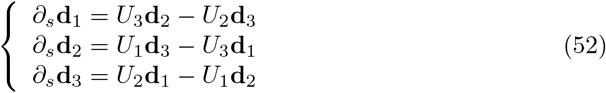

which is derived from (3). We can then recover the centerline **r**^*a*^ by integrating *∂*_*s*_**r**^*a*^ = **d**_3_. The PFR centerline **r**^*p*^ and the orthonormal frame **g**_1_, **g**_2_, and **g**_3_ follow from (9) and (10).

## C Numerics

We define the nondimensional variables for arc length *x*, time *y*, strains *u*_*i*_, shear forces *h*_*i*_, and concentrated shear forces 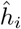 as follows

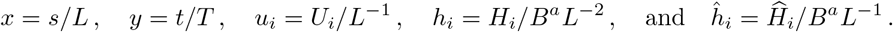

The equilibrium equations (12), with (21) and (23), are solved by seeking for a minimizer of the energy (26) in the adimensional form

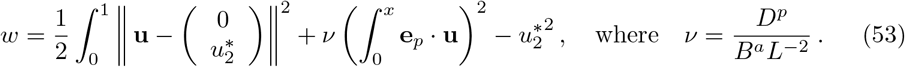

In the formula above **u** = (*u*_1_, *u*_2_) is the adimensional bending vector and 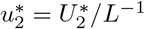 is the adimensional target bending. We find the minimizer using the gradient descend method

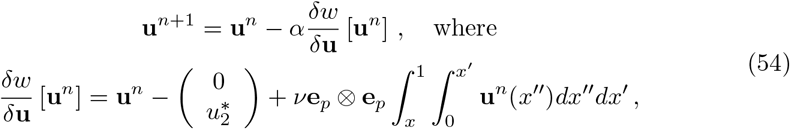

and *α* is a conveniently chosen step size. We initiate the algorithm with 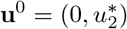, and then we iterate (54) until ‖**u**^*n*^+1 − **u**^*n*^‖ falls below a pre-set tolerance parameter.

The dynamic equations (17) are recast, and then solved, in terms of the shear vector *γ* = (*γ*_1_, *γ*_2_) where

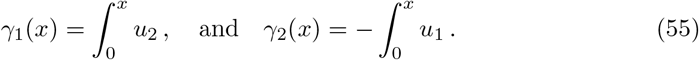

After solving for *γ*, we obtain the bending strains by differentiation. The equation for *γ* is

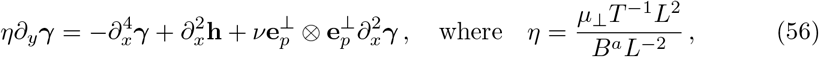

and **h** = (*h*_1_, *h*_2_). The corresponding boundary conditions are given by

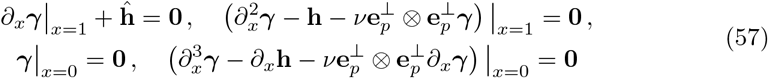

where 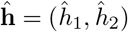. The above (57) are point-wise conditions, which do not involve integral terms as in (19). Avoiding this non-locality allows for an easier numerical implementation. The finite difference scheme we employ to solve (56)-(57) is illustrated in the remainder.

We consider the discrete time sequence *y*_*n*_ = *n*Δ*y* with *n* = 0, 1, 2, . . . and we define *γ*^*n*^(*x*) = *γ*(*y*_*n*_, *x*), **h**^*n*^(*x*) = **h**(*y*_*n*_, *x*), and 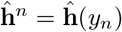. Equation (56) is discretized in time with the one-step (semi-implicit) numerical scheme

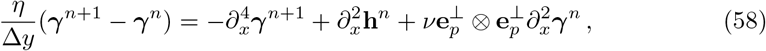

and complemented by the boundary conditions

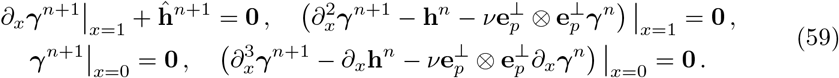

The adimensional arc length interval [0, 1] is discretized uniformly in *M* + 1 points *x*_*k*_ = *k*Δ*x*, with *k* = 0, 1, . . . , *M* = 1/Δ*x*. We also consider the extra “ghost points” *x*_*k*_ = *k*Δ*x* with *k* = −1, *M* + 1, *M* + 2. The discrete values 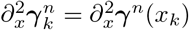 of the second derivative in (58) are approximated by the finite difference scheme

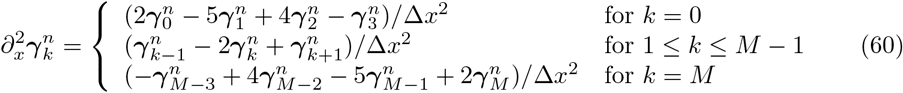

where 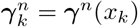. Analogous formulas are employed for the second derivative of **h**^*n*^. The discrete values 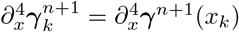 of the forth derivative in (58) are given by the scheme

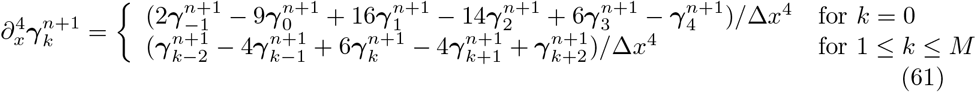

which involves the ghost points values *γ*_*−1*_, *γ*_*M* +1_, and *γ*_*M* +2_. The discretized approximations of the boundary conditions (59) are given by

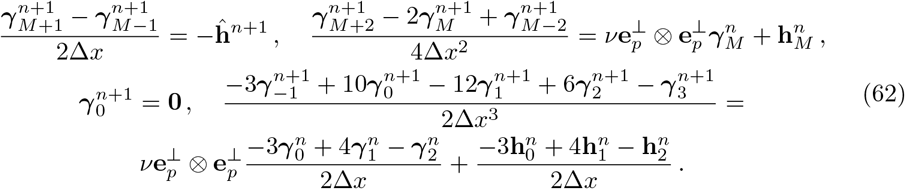

The previous formulas give us the expressions for the ghost points’ values 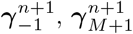, and 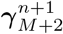 in terms of 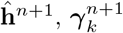, and 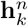 with 0 ≤ *k* ≤ *M*. These expressions are then plugged in (61). In turn, the iterative scheme (58) allows to calculate 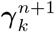 from 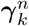 with 0 ≤ *k* ≤ *M*, while incorporating the boundary conditions (59) in the numerical solution.

The scheme is iterated for several time periods until a periodic solution is reached.

## D Estimation of dynein forces and mechanical parameters

We obtain the history of shear forces presented in Figure 7 by solving the following inverse dynamical problem. We assign first a history of normalized bending strains (0, *u*_2_(*x, y*)), periodic in time, that imitates the experimentally observed Chlamydomonas flagellar curvatures reported in [23]. We use the following model

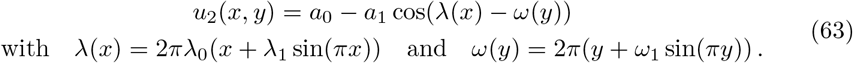

Then, we calculate the shear forces that generate said history of bending strains for an Ax beating in a viscous fluid (without extra-axonemal structures attached to it).

Equation (56) with *ν* = 0 and 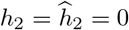 defines exactly the dynamics of this system. We can solve for *h*_1_ and 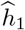 explicitly, obtaining 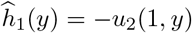 and

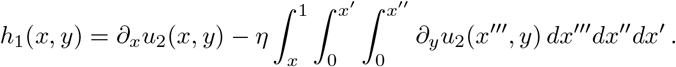

In the dynamic simulation in Figure 7 we used the following numeric values for the physical parameters of the system. The bending modulus of the Ax is *B*^*a*^ = 840 pN · *μ*m^2^, taken from [37]. We set *L* = 28 *μ*m, *T* = 25 ms, and *μ*_⊥_ = 3.1 fN · *s · μ*m^−2^, which are all values estimated in [25]. The angle between spontaneous bending plane and the Ax-PFR joining line *ϕ*^*p*^ = −2*π*/9 is estimated from micrographs in [20] and [4]. Without direct measurements for *D*^*p*^, we set *ν* = 20 as the value giving the most balanced mechanical interplay between Ax and PFR (the value of *ν* with the most evenly spread-out bending vector’s solutions in Figure 6). For the bending strains parameters in (63) we took *a*_0_ = 7.8, *a*_1_ = 7.5, *λ*_0_ = 1.85, *λ*_1_ = 0.1, and *ω*_1_ = −0.1.

